# Genetic differences between 129S substrains affect antiretroviral immune responses

**DOI:** 10.1101/2022.08.10.503554

**Authors:** Robert Z. Zhang, Vincent Mele, Lia Robben, Melissa Kane

## Abstract

Inbred mouse lines vary in their ability to mount protective antiretroviral immune responses, and even closely related strains can exhibit opposing phenotypes upon retroviral infection. Here, we found that 129S mice inherit a previously unknown mechanism for the production of anti-murine leukemia virus (MLV) antibodies and control of infection. The resistant phenotype is controlled by two dominant loci that are independent from known MLV-resistance genes. We also show that production of anti-MLV antibodies in 129S7, but not 129S1 mice is independent of interferon gamma (IFNγ) signaling. Thus, our data indicate that 129S mice inherit an unknown mechanism for control of MLV infection and demonstrate that there is genetic variability in 129S substrains that affects their ability to mount antiviral immune responses.

**Importance:** Understanding the genetic basis for production of protective antiviral immune responses is crucial for the development of novel vaccines and adjuvants. Additionally, characterizing the genetic and phenotypic variability in inbred mice has implications for the selection of strains for targeted mutagenesis, choice of controls, and for broader understanding of the requirements for protective immunity.

## Introduction

Resistance and sensitivity to viral infections depends greatly upon the genetic make-up of the host. Model organisms such as inbred mice have proved to be invaluable for the investigation of the pathways underlying protective immune responses due to the inability to perform genetic manipulations in humans. In this regard, mouse models of murine retroviruses, such as murine leukemia virus (MLV) and mouse mammary tumor virus (MMTV) have provided essential insights into the molecular mechanisms underlying anti-viral immune responses. MLV is transmitted as an exogenous virus through the blood and milk or as an endogenous stably integrated provirus and primarily infects cells of lymphoid origin (1). Pathogenic MLVs are often a mixture of ecotropic and polytropic [mink cell focus forming (MCF)] viruses, and some contain an additional replication-defective spleen focus forming virus (SFFV) component that promotes pathogenesis (1). These viruses cause a range of diseases in susceptible mice, including lymphomas, leukemias, rhabdomyosarcoma, neurological disorders, and immunosuppression (2-6). Mice from retrovirus-susceptible strains, such as BALB/c mice, sense retroviral pathogens, as indicated by the fact that they initiate an antiretroviral response. However, this response is not long-lasting and is unsuccessful in controlling virus replication (7, 8), which is most likely due to the numerous mechanisms of immune evasion employed by retroviruses (9-13). In contrast, pathogen detection in resistant mice translates into a robust, long-lasting, and virus-neutralizing immune response comprised of both T and B cells responses (7, 9, 14, 15).

Classical genetics approaches in mice have identified several genes that play a role in directly restricting MLV replication, or in the generation of protective antiviral immune responses [Table 1 and recently reviewed in (16)]. *Fv1* is a capsid-specific restriction factor encoded by an endogenous retroviral *Gag* gene that governs tropism of MLV. Alleles of *Fv1* are defined by their ability to block specific subclasses of MLV, with incompatibility resulting in a post-entry block to infection, (17-20) and in some cases, stimulation of cross-protective neutralizing adaptive immune responses (21). *Fv2* is a dominant Friend MLV (FV) susceptibility gene that facilitates splenomegaly and erythroleukemia induction by SFFV by promoting erythropoietin receptor signaling (22, 23). *Rfv3* is a dominant FV resistance gene that is encoded by *Apobec3*. The resistant allele of *Apobec3* inherited by C57BL/6 and C57BL/10 mice restricts FV replication and promotes neutralizing antibody (Ab) responses (24). *Vic1* is a recessive MLV and MMTV resistance gene that is encoded by *H2-Ob*, the beta subunit (Oβ in mice and DOβ in humans) of the nonclassical major histocompatibility complex class II (MHC-II)-like molecule H2-O (HLA-DO or DO in humans) (25). The resistant, functionally null allele of *vic1* inherited by I/LnJ mice results in sustained production of virus-neutralizing Abs. Transfer of the I/LnJ *vic1* locus to multiple retrovirus susceptible backgrounds, which contain a functional allele of *Ob*, such as BALB/cJ (BALB/c^v*ic1I/LnJ*^ congenic mice) confers the ability to produce neutralizing Abs against both MMTV and MLV (25, 26). Finally, adaptive immune responses against retroviruses in mice (and humans) are also affected by classical MHC class I and II genes, with specific haplotypes directing virus-specific cytotoxic T cell and Ab responses (16, 22).

**Table 1.**
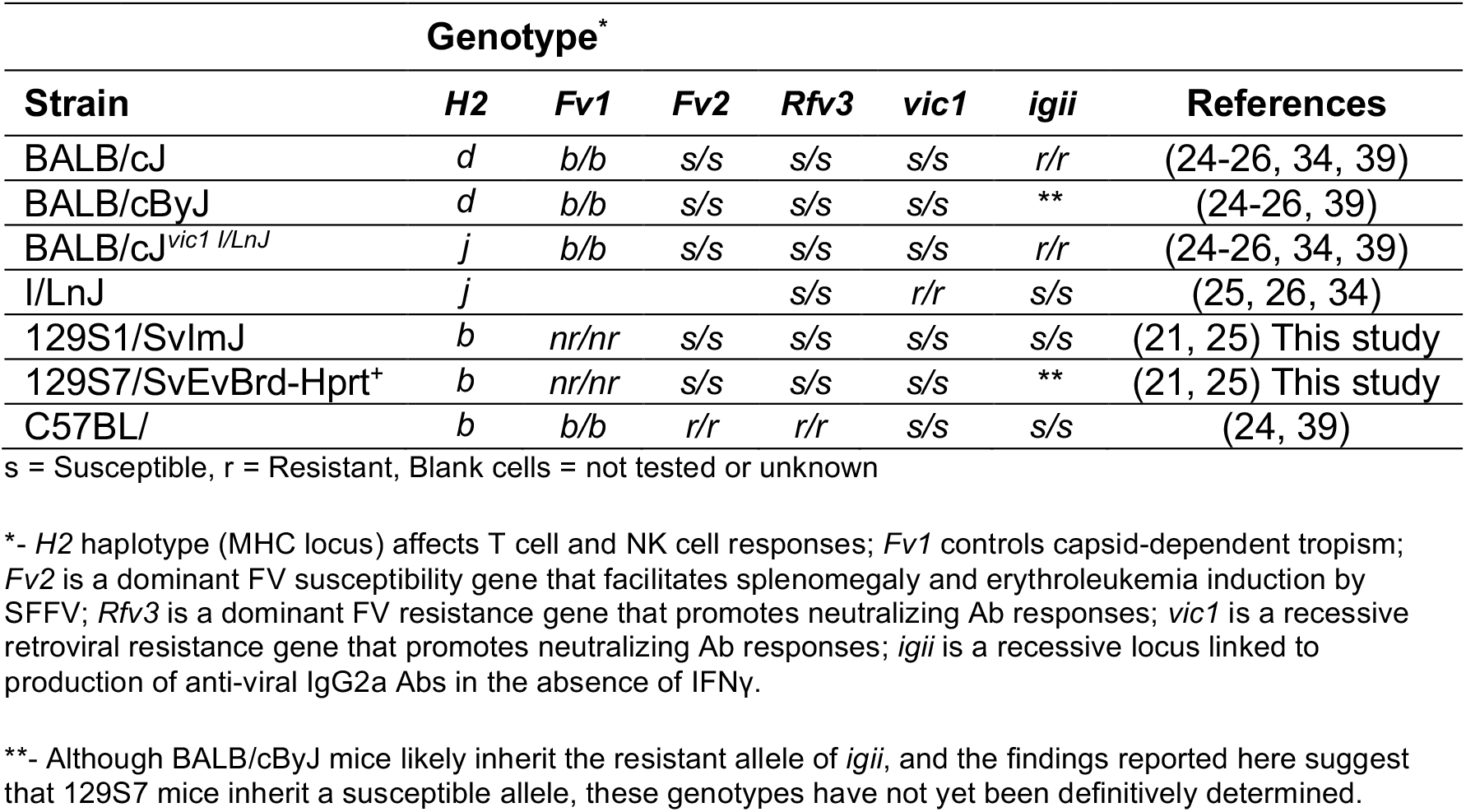
Genotypes of inbred strains at MLV-resistance loci

| Genotype* |  |  |  |  |  |  |  |
| --- | --- | --- | --- | --- | --- | --- | --- |
| Strain | H2 | Fv1 | Fv2 | Rfv3 | vic1 | igii | References |
| BALB/cJ | d | b/b | s/s | s/s | s/s | r/r | (24-26, 34, 39) |
| BALB/cByJ | d | b/b | s/s | s/s | s/s | ** | (24-26, 39) |
| BALB/cJ<sup>vic1 I/LnJ</sup> | j | b/b | s/s | s/s | s/s | r/r | (24-26, 34, 39) |
| I/LnJ | j |  |  | s/s | r/r | s/s | (25, 26, 34) |
| 129S1/SvImJ | b | nr/nr | s/s | s/s | s/s | s/s | (21, 25) This study |
| 129S7/SvEvBrd-Hprt<sup>+</sup> | b | nr/nr | s/s | s/s | s/s | ** | (21, 25) This study |
| C57BL/ | b | b/b | r/r | r/r | s/s | s/s | (24, 39) |

s = Susceptible, r = Resistant, Blank cells = not tested or unknown
\*- *H2* haplotype (MHC locus) affects T cell and NK cell responses; *Fv1* controls capsid-dependent tropism; *Fv2* is a dominant FV susceptibility gene that facilitates splenomegaly and erythroleukemia induction by SFFV; *Rfv3* is a dominant FV resistance gene that promotes neutralizing Ab responses; *vic1* is a recessive retroviral resistance gene that promotes neutralizing Ab responses; *igii* is a recessive locus linked to production of anti-viral IgG2a Abs in the absence of IFNγ .
\*\* - Although BALB/cByJ mice likely inherit the resistant allele of *igii*, and the findings reported here suggest that 129S7 mice inherit a susceptible allele, these genotypes have not yet been definitively determined.

Ab responses to soluble protein and carbohydrate antigens in mice are generally restricted to the IgG1 and IgG3 isotypes (27, 28), while infection of mice with a variety of viral and parasitic pathogens results in a unique bias in class switching to IgG2a (or IgG2c) Abs (29-31), including antiretroviral Ab responses in I/LnJ mice (7). In mice, the Th1-type cytokine interferon gamma (IFNγ) has been well established as the canonical signal for class switching to the IgG2a/c isotype both in vitro and in the context of infection (32). Specifically, in B6 mice, IFNγ is required for Ab responses to MLV infection and for long-term viral control (33), and in I/LnJ mice, IFNγ is essential to produce antibodies of any isotype against retroviral infections (7, 26). However, we recently reported that in BALB/cJ^*vic1i/i*^ mice, IFNγ is dispensable for the production of anti-MLV IgG2a Abs (34), indicating that BALB/cJ mice inherit an alternative mechanism for stimulating antiviral IgG2a Ab production upon viral infection. The ability to generate IFNγ-independent Ab responses in BALB/cJ mice is controlled by a single recessive locus on Chromosome (Chr) 9, which we have named *i*nterferon-*g*amma *i*ndependent *I*gG2a (*igii*) (34).

Reverse genetics approaches in mice are a powerful tool to understand the function of specific genes and have provided essential research models for numerous diseases. Prior to the development of CRISPR technology, which allowed for targeted mutagenesis on multiple genetic backgrounds, most ES cell lines used for targeted mutagenesis came from substrains of 129 mice. The 129 lineage has a complex history, with multiple outcrossing and bottleneck events, resulting in three major lineages (Parental, Steel, and *Ter*), with substantial substrain variability ((35, 36) and Figure S1). This complex history has important implications for investigations using 129 mice, and on analysis of targeted mutations that have been crossed to other genetic backgrounds. Here, we investigated anti-MLV immune responses in 129S substrains and determined that 129S1 mice inherit two loci that control antiviral immunity. We also demonstrate that genetic differences between 129S substrains affect IFNγ-independent antiviral immune responses.

## Results

### A dominant genetic mechanism controls anti-MLV immune responses in 129S1 mice

Previous reports demonstrated that 129P2 mice (representative of the Parental 129 substrains) inherit the *Fv1*^nr^ allele, which restricts B-tropic and some N-tropic, but not NB-tropic strains of MLV (21, 37, 38). 129P2 infected with a B-tropic FV produce robust cross-protective neutralizing Ab responses due to this *Fv1*-incompatibility. However, although they produce FV-specific Abs upon infection with an NB-tropic FV, since they inherit susceptible alleles of *Fv2* and *Rfv3*, they do not control the infection (21, 39). These reports additionally suggest that the entire 129 lineage is *Fv1*^nr/nr^ and *Fv2*^s/s^ (21, 23, 37, 38). Furthermore, genotyping of two Steel substrains (see Materials and Methods) at the *Rfv3* locus indicates that the 129 lineage is *Rfv3*^*s/s*^. To investigate antiviral responses in 129 Steel substrains, we infected 129S1/SvImJ (129S1; developed as a control for steel-derived ES cells (40)) mice with Rauscher-like-MLV (RL-MLV) and monitored them for antiviral Ab production, development of splenomegaly, and viral titers in the spleen. RL-MLV is a mixture of a NB-tropic ecotropic virus and a polytropic MCF virus that causes erythroid leukemias; the mixture does not contain an SFFV component (3), therefore the *Fv2* allele does not affect disease outcome. While susceptible control BALB/cByJ (ByJ) mice do not produce a robust Ab response and were unable to control infection, 129S1 mice produced a robust IgG2a Ab response against RL-MLV and the majority (83%) cleared infection and did not develop splenomegaly (Figures 1 and S2). To determine whether the ability to produce protective anti-MLV Ab responses is inherited in a dominant or recessive fashion, we crossed 129S1 mice to BALB/cByJ mice (Figure 2A). F_1_ mice generated from these crosses were infected with RL-MLV and screened for antiviral Abs. F_1_ mice from crosses of both directions [(129S1xByJ) and (ByJx129S1) produced antiviral Abs, although IgG2a productions was slightly lower in (ByJx129S1) F_1_ animals (Figure 1A). Additionally, the majority of F_1_ mice eliminated the virus and did not develop splenomegaly (Figure 1B-C), indicating that resistance to RL-MLV is inherited in a dominant fashion.

**Figure 1.**
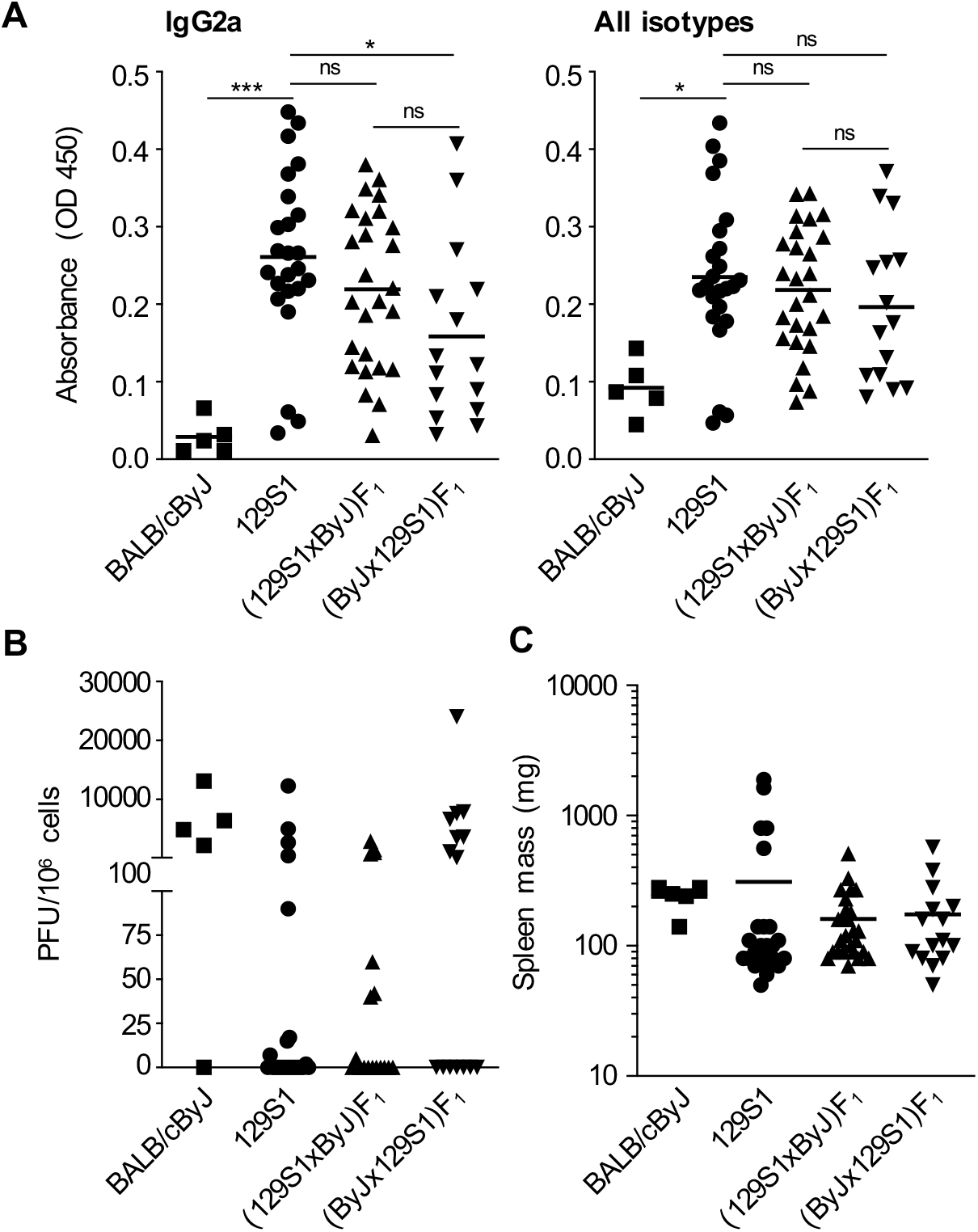
Anti MLV-Ab production in 129S1 mice is controlled by a dominant genetic mechanism. **A)** BALB/cByJ, 129S1, or F_1_ mice were infected with RL-MLV and monitored for IgG2a-specific antibodies (left) or total Igs (right) against RL-MLV virion proteins by ELISA 8-10 weeks post infection. ns, not significant; *, p<0.05; ***, p<0.001 **B)** Spleen cells from RL-MLV infected mice were subjected to an infectious center assay 8-10 weeks post infection. **C)** Spleen weights of RL-MLV infected mice at 8-10 weeks post infection.

**Figure 2.**
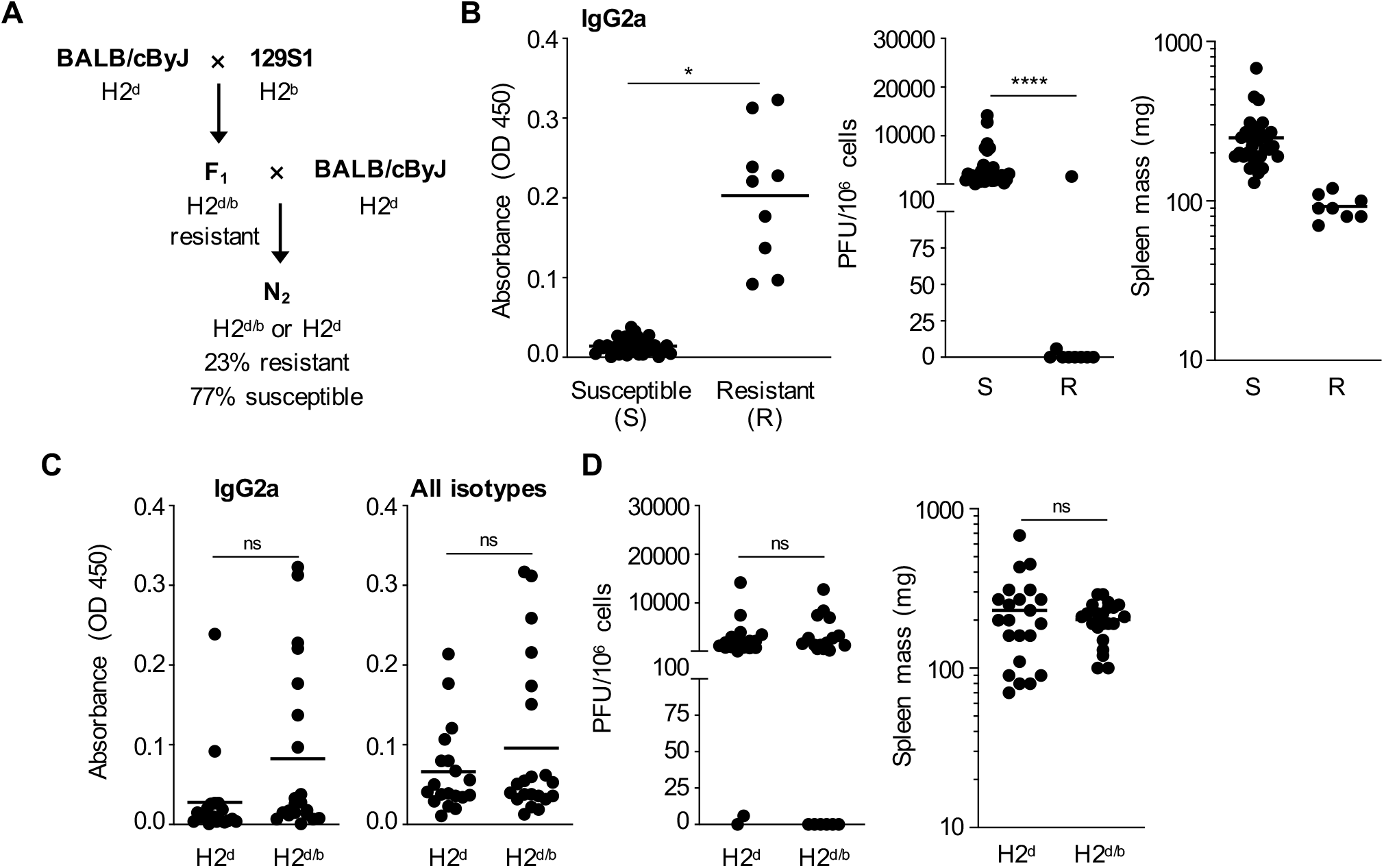
Anti MLV-Ab production in 129S1 mice is controlled by two non-MHC loci. **A)** Diagram of breeding scheme used to determine Mendelian inheritance of the mechanism controlling IFNγ-independent IgG2a Ab production. F_1_ and N_2_ crosses were conducted in both directions, with just one direction shown here for simplicity. **B)** N_2_ mice generated from crossing (129S1xByJ)F_1_ and ByJ mice as shown in A) were infected with RL-MLV. Eight weeks post infection, their sera were tested for IgG2a Abs against MLV virion proteins by ELISA (left), mice were classified as “resistant” if their sera reacted with MLV virion proteins with an absorbance greater than 0.1 at OD 450 with an IgG2a-specific secondary. Susceptible and resistant mice were then tested for presence of infectious virions in their spleens by XC plaque assay (center), and spleen weights were measured (right). **C)** MHC haplotypes of N_2_ mice from B) were determined by FACS analysis and then levels of IgG2a-specific antibodies (left) or total Igs (right) reactive against RL-MLV virion proteins by ELISA were compared. **D)** Presence of infectious virions in spleen (left) and spleen weights (right) of N_2_ mice grouped by MHC haplotype.

### Two loci control anti-MLV Ab responses in 129S1 mice

We next investigated how many loci control anti-MLV Ab production by crossing resistant (129S1xByJ) F_1_ mice to susceptible ByJ mice to generate N_2_ mice (Figure 2A). We found that 9 out of 40 (23%) N_2_ mice produced antiviral IgG2a Abs (Figure 2B). We then determined the viral titers in the spleen and spleen weights of Ab negative and Ab positive N_2_ mice and found that the Ab positive (resistant), but not the Ab negative (susceptible) N_2_ mice cleared the infection and had normal spleen weights (Figure 2B). This three:one ratio of susceptible to resistant N_2_ animals indicates that two loci control the production of protective anti-RL-MLV Ab responses in 129S1 mice. Since the major histocompatibility complex (MHC) locus is a major factor in controlling antiviral immunity, including anti-MLV responses (16, 22), and 129S1 mice inherit the protective H2^b^ haplotype while ByJ mice inherit the susceptible H2^d^ haplotype (Table 1), we next investigated whether the MHC locus is one of the genetic determinants for anti-MLV Ab responses in 129S1 mice. MHC haplotypes of N_2_ mice were determined by staining peripheral blood lymphocytes with haplotype-specific Abs for MHC Class I (H2-D^b^, H2-D^d^) and Class II (I-A^b^, I-A^d^). We did not observe significant differences in Ab production, viral titers, and spleen weights in RL-MLV infected H2^b/d^ and H2^d/d^ N_2_ mice (Figure 2C-D and Table S1), indicating that control of infection in N_2_ mice is not determined by inheritance of the resistant H2^b^ haplotype and that non-MHC loci control antiviral responses in 129S1 mice.

### IFNγ-signaling is dispensable for anti-MLV IgG2a-specific Ab production in 129S7 mice

Following our previous finding that BALB/cJ mice congenic for the resistant allele of *vic1* produce antiviral IgG2a Abs in the absence of IFNγ (34), we wondered whether this pathway was unique to this genetic background. Earlier reports indicated that 129S mice also inherit a pathway for IFNγ-independent production, as IFNγ-receptor 1 (IFNγR)-deficient 129S7 (G129) mice produce IgG2a Abs against lymphocytic choriomeningitis virus (LCMV) (41) and (lactate dehydrogenase-elevating virus) LV (42). We therefore investigated whether IFNγR^-/-^ 129S7 mice produce IFNγ-independent IgG2a Abs against RL-MLV. Control IFNγR^-/-^ ByJ (which do not inherit the resistant *vic1* allele, Table 1) mice did not produce Abs of any isotype and did not control infection, while IFNγR^-/-^ 129S7 mice produced IgG2a Abs against RL-MLV (Figure 3A and Figure S2). Although Ab production was lower than in IFNγR-sufficient 129S1 mice, the majority of IFNγR^-/-^ 129S7 mice (60%) cleared infection and did not develop splenomegaly (Figure 3 and Table 1). Therefore, like BALB/cJ mice, 129S7 mice inherit a mechanism for noncanonical, IFNγ-independent antiretroviral Ab production.

**Figure 3.**
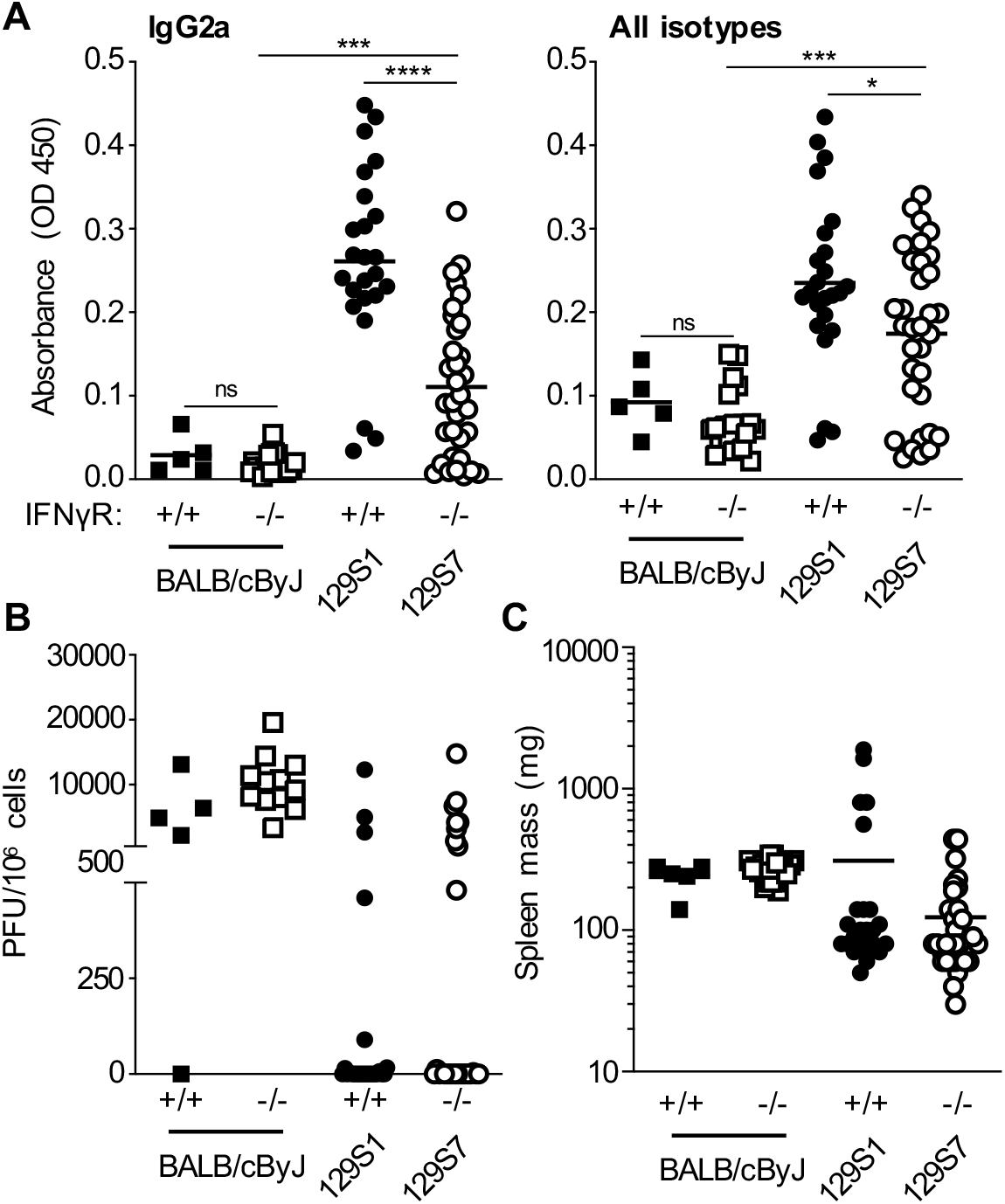
IFNγR-independent anti-MLV IgG2a Ab production in 129S7 mice. **A)** Mice of the indicated genotypes were infected with RL-MLV and monitored for IgG2a-specific antibodies (left) or total Igs (right) against RL-MLV virion proteins by ELISA 8-10 weeks post infection. ns, not significant; *, p<0.05; ***, p<0.001; ****, p<0.0001 **B)** Spleen cells from RL-MLV infected mice were subjected to an infectious center assay 8-10 weeks post infection. **C)** Spleen weights of RL-MLV infected mice at 8-10 weeks post infection.

### IFNγ-independent anti-MLV IgG2a production in 129S7 mice is a complex trait

We next sought to investigate whether IFNγ-independent antiretroviral Ab production in 129S7 mice and BALB/cJ mice is controlled by the same genetic mechanism. Since the resistant allele of *igii*-inherited by BALB/cJ mice is recessive, we reasoned that IFNγR^-/-^ F_1_ animals inheriting the 129S7 and BALB/c allele would only produce antiviral IgG2a Abs if the 129S7 allele of *igii* is resistant. IFNγR-deficient mice are available on the BALB/cByJ background, and we crossed these mice to IFNγR^-/-^ 129S7 mice. F_1_ mice generated from these crosses were infected with RL-MLV and screened for antiviral Abs. These F_1_ progeny did not display a clear phenotype, with ∼30% of mice producing Abs and controlling infection while the majority did not produce Abs, developed splenomegaly, and had infectious virus in their spleens (Figure 4 and Table S1). Therefore, the phenotype is not fully penetrant, and the genetic basis for IFNγ-independent antiretroviral Ab production in 129S7 mice cannot be determined from these crosses. Furthermore, although these experiments suggest that 129S7 mice do not inherit a resistant allele of *igii*, BALB/cByJ and BALB/cJ mice have been separated since 1935, and *vic1* congenics are not available on the ByJ background, therefore we cannot exclude the possibility that ByJ and BALB/cJ mice inherit different alleles of *igii*.

**Figure 4.**
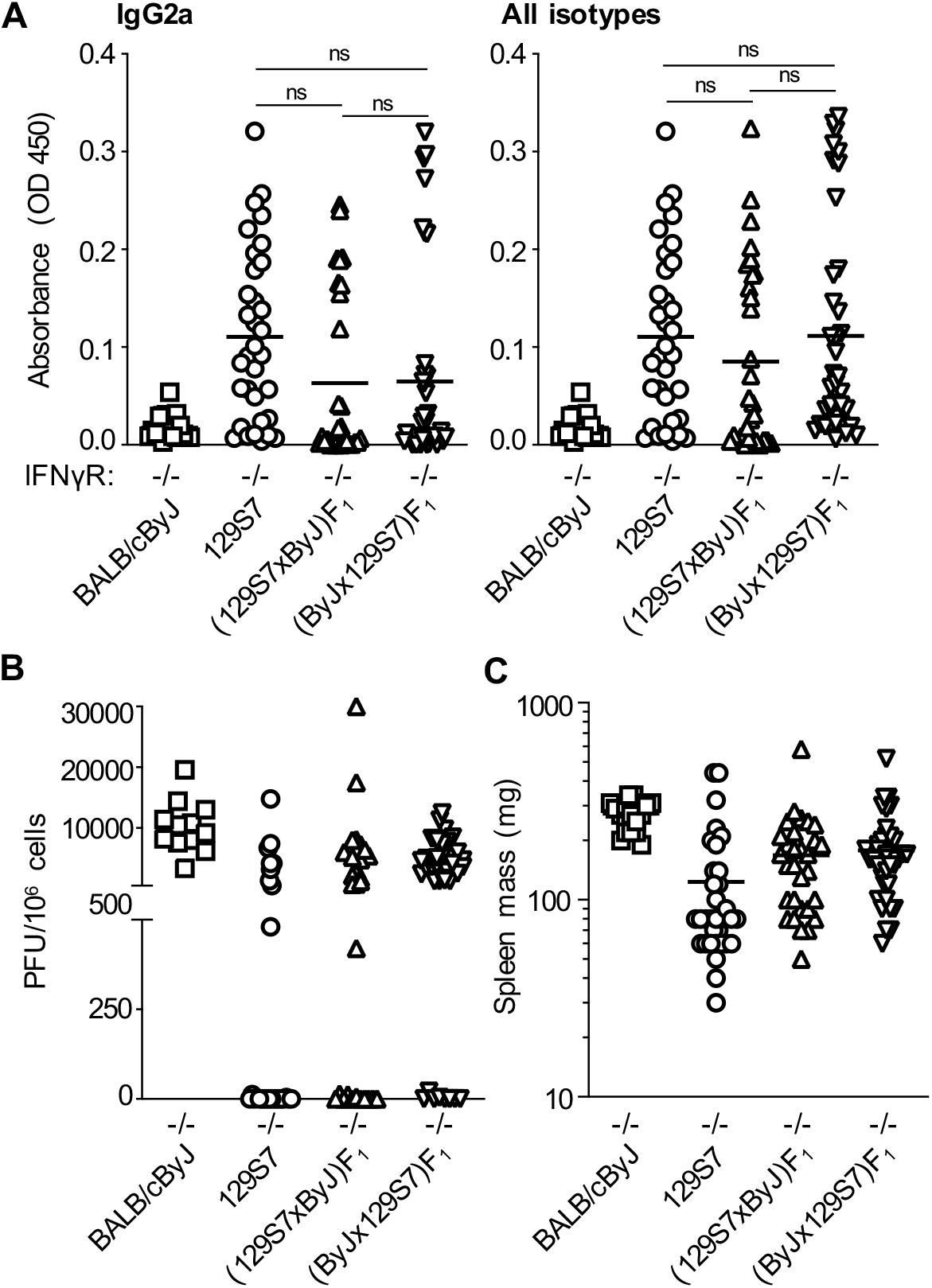
IFNγR-independent anti-MLV Ab production in 129S7 mice is a complex trait. **A)** IFNγR^-/-^ BALB/cByJ, 129S7, and F_1_ mice of the indicated genotypes were infected with RL-MLV and monitored for IgG2a-specific antibodies (left) or total Igs (right) against RL-MLV virion proteins by ELISA eight weeks post infection. ns, not significant **B)** Spleen cells from RL-MLV infected mice were subjected to an infectious center assay eight weeks post infection. **C)** Spleen weights of RL-MLV infected mice at eight weeks post infection.

### Anti-MLV Ab production in 129S1 mice is IFNγ-dependent

Since we were unable to determine the genetic basis for IFNγ-independent antiretroviral Ab production in 129S7 mice by crossing to IFNγR^-/-^ ByJ mice, and these strains are deficient for the IFNγ receptor rather than the cytokine, we decided to generate IFNγ-deficient 129S mice for the investigation of IFNγ-independent Ab responses. Considering that wild-type mice of the precise genetic background of 129S7-*Ifngr*^*-/-*^ no longer exist, we selected the 129S1 background for the knock-out of IFNγ, as they are the control strain recommended by the Jackson Laboratory for lines derived from multiple 129S substrains. We used a previously validated CRISPR/Cas9 approach that targeted introns 1 and 3 of the *Ifng* gene, thereby deleting exons 2 and 3, removing four of six alpha helices ((43) and Figure S3A). We infected heterozygous control and IFNγ^-/-^ mice from two founder lines with RL-MLV and monitored them for antiviral Ab production. Although IFNγ sufficient controls produced antiviral IgG2a Abs and cleared the infection, IFNγ^-/-^ 129S1 mice from either founder line failed to produce a robust antiviral Ab response of any isotype (Figure 5). Additionally, less than 20% of IFNγ^-/-^ 129S1 mice cleared the infection, and the majority developed splenomegaly (Figure 5B-C and Table S1), indicating that unlike 129S7 mice, 129S1 mice do not inherit a pathway for directing IFNγ-independent antiretroviral Ab responses.

**Figure 5.**
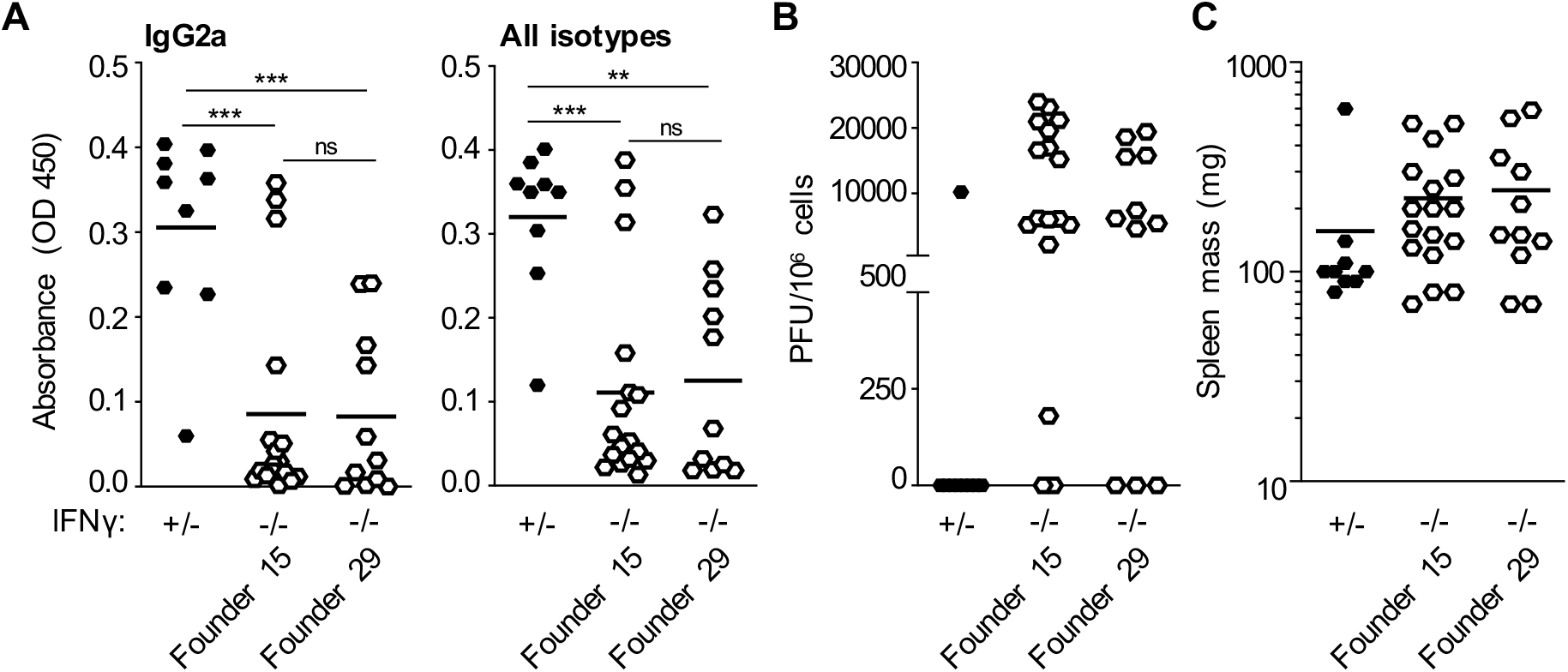
Anti-MLV Ab responses in 129S1 mice are IFNγ-dependent. **A)** IFNγ-deficient and heterozygous control mice from two 129S1 founder lines were infected with RL-MLV and monitored for IgG2a-specific antibodies (left) or total Igs (right) against RL-MLV virion proteins by ELISA eight weeks post infection. ns, not significant; **, p<0.01; ***, p<0.001 **B)** Spleen cells from RL-MLV infected mice were subjected to an infectious center assay eight weeks post infection. **C)** Spleen weights of RL-MLV infected mice at eight weeks post infection.

### Genetic differences, not alternative receptor usage, underlie differential requirements for IFNγ-signaling in 129S mice

To determine if antiviral IgG2a Ab responses in 129S7 mice are the result of IFNγ signaling through an alternative receptor or result from genetic differences between 129S7 and 129S1 mice, we crossed IFNγR^-/-^ 129S7 mice to IFNγ^-/-^ 129S1 mice and then intercrossed the resulting F_1_ generation (Figure 6A). If IFNγ signaling through an alternative receptor explained virus resistance in IFNγR^-/-^ 129S7 mice, we would expect to observe antiviral Ab production in IFNγR^-/-^, but not IFNγ^-/-^, or IFNγ/IFNγR^-/-^ F_2_ mice. On the other hand, if 129S7 inherit an alternative pathway for IFNγ-independent antiviral Ab production, we would expect a mixture of resistant and susceptible F_2_ mice independently of inheritance of IFNγR- or IFNγ-deficiency. As expected, F_2_ mice sufficient for both cytokine and receptor produced anti-MLV IgG2a-specific Abs and cleared the infection (Fig 6B-C). However, antiviral Ab production was not observed in the majority of IFNγR^-/-^ F_2_ mice, and only 10% cleared the infection (Figure 6B-C). Interestingly, more IFNγ^-/-^ and IFNγ/IFNγR^-/-^ F_2_ mice produced anti-MLV Abs and cleared infection than IFNγR^-/-^ F_2_ mice, and 64% of IFNγ^-/-^ and 100% of IFNγ/IFNγR^-/-^ F_2_ mice cleared the infection (although only four double-deficient mice were infected) (Figure 6B-D and Table S1). These data strongly suggest that IFNγ does not signal through an alternative receptor to stimulate antiviral Abs in 129S7 mice and that an unknown genetic factor(s) present in 129S7, but not 129S1 mice controls IFNγ-independent antiretroviral Ab responses.

**Figure 6.**
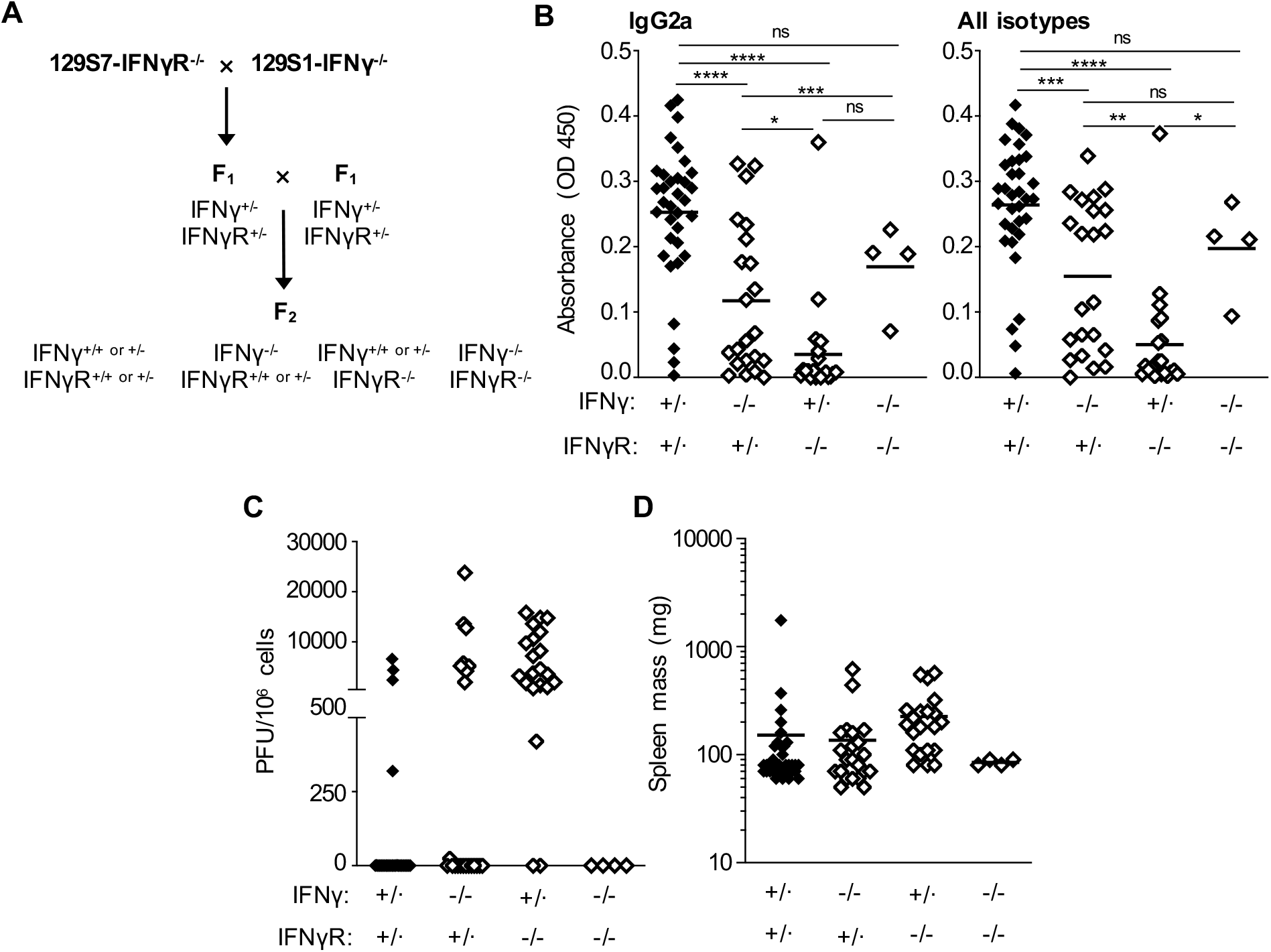
Genetic differences between 129S substrains affect IFNγ-independent anti-MLV Ab responses. **A)** Diagram of breeding scheme used to generate IFNγ, IFNγR, and IFNγ/IFNγR-deficient mice of mixed 129S7 and 129S1 genetic background. **B)** F_2_ mice of the indicated genotypes generated from intercrossing (129S7-*Ifngr1*^-/-^ x 129S1-*Ifng*^-/-^)F_1_ mice as shown in A) were infected with RL-MLV. 8-10 weeks post infection, their sera were tested for IgG2a-specific antibodies (left) or total Igs (right) against RL-MLV virion proteins by ELISA. +/·, either heterozygous or homozygous wild type. ns, not significant; *, p<0.05; **, p<0.01; ***, p<0.001; ****, p<0.0001 **C)** Spleen cells from RL-MLV infected F_2_ mice were subjected to an infectious center assay eight weeks post infection. **D)** Spleen weights of RL-MLV infected F_2_ mice at eight weeks post infection.

## Discussion

Here, we demonstrate that 129S1 mice inherit a previously unknown mechanism that controls the production of protective antiviral Ab production upon MLV infection. This phenotype is inherited in a dominant fashion and is controlled by two loci (Figures 1-2). Importantly, these loci are distinct from previously characterized genes that restrict retroviral replication (*Fv1, Fv2, Rfv3*) or promote antiviral immune responses (*vic1*, MHC locus, *Rfv3*), indicating that the pathway in 129S1 mice potentially encodes for a novel mechanism for the stimulation of protective immunity. Unlike I/LnJ and C57BL/6 mice, which fully clear RL-MLV infection (15, 26), MLV resistance in 129S1 mice is not fully penetrant, as ∼15% of infected mice fail to produce antiviral Abs, develop splenomegaly, and retain infectious virus in their spleens (Figures 1 and 5, Table S1). Additionally, although this mechanism does not appear to be sex-linked, there was a reduction in IgG2a Ab production in (ByJx129S1) F_1_ mice, and fewer of them cleared the infection than 129S1 or (129S1xByJ) F_1_ mice (Figure 1 and Table S1), indicating that epigenetic or environmental factors may play a role in controlling anti-MLV responses in 129S1 mice.

IFNγ is a well-established signal for stimulating CSR to the IgG2a isotype both *in vitro* and in the context of various viral infections (21, 37, 38), including FV infection in B6 mice (22) and RL-MLV and MMTV infection in I/LnJ mice (6, 15). We previously demonstrated that BALB/cJ mice inherit a pathway for the generation of IFNγ-independent IgG2a antiviral Ab responses that is controlled by a single recessive locus, *igii*, which is mapped to Chr. 9 (34). Here, we demonstrate that 129S7 mice also inherit a pathway for IFNγ-independent anti-MLV Ab production. However, the mechanism for IFNγR-independent anti-RL-MLV responses in 129S7 mice is a complex genetic trait, and we were unable to determine whether the pathway is controlled by *igii* or another locus (or multiple loci). Nevertheless, the ability of both BALB/cJ and 129S7 mice to produce IFNγ-independent Abs against a variety of infections including retroviruses, herpes simplex virus (HSV), LCMV, LV, and parasitic infections (34, 41, 42), indicates that alternative pathways in addition to the canonical IFNγ-dependent pathway can stimulate neutralizing Ab responses. Since production of IFNγ-independent Ab responses in BALB/cJ mice is independent of *vic1* for HSV and VSV infections (but not MLV infections), this indicates that the ability to produce anti-viral Ab responses and the ability to produce these responses in the absence of IFNγ signaling are controlled by distinct genetic mechanisms. Although we anticipate this is also the case for 129S7 mice, it cannot be definitively demonstrated. These findings additionally highlight the importance of utilizing mice of multiple genetic backgrounds to investigate the requirements for stimulating protective antiviral immunity.

Following decades of widespread use of 129 ES cell lines, two reports from 1997 investigated the variation between 129 substrains and found substantial genetic variation resulting from genetic drift, genetic contamination, and residual heterozygosity from backcrossing programs (35, 36). This initiated efforts to clarify the nomenclature for the improvement of gene targeting experiments and the selection and consideration of controls for comparing phenotypes of genes targeted in ES cells of different 129 substrains (35, 36, 44). These findings further suggested important features that should be considered for targeted mutagenesis using 129 substrains 1) chimeras produced using 129 ES cells should be crossed to a genetically matched substrain for maintenance on the 129 background and 2) selection of a 129 substrain for targeted mutagenesis should involve consideration of different genetic traits between substrains (35). Like many mice generated prior to these recommendations, chimeric IFNγR^-/-^ 129S mice were not crossed to a genetically matched substrain, and the appropriate wild-type control strain is unavailable. The 129S1 substrain is therefore recommended as the control inbred strain for steel substrain-derived ES cell lines (40). However, we found that while 129S7 mice produce antiviral Ab responses in the absence of IFNγ, this mechanism is not inherited by 129S1 mice (Figure 5). Some reports have suggested that IFNγ can signal through an alternative receptor in the absence of *Ifngr1* (45-47), but this does not explain the differences between 129S1 and 129S7 substrains reported here, as IFNγR^-/-^ (129S1×129S7) F_2_ mice did not produce antiviral Abs (Figure 6). Therefore, for the purposes of investigating the requirements for IFNγ in antiviral defenses, 129S1 mice are not an appropriate control for 129S7-*Ifngr*^*-/-*^ mice. The higher incidence of IFNγ-independent Ab production in IFNγ^-/-^ and IFNγ/IFNγR^-/-^ F_2_ mice than IFNγR^-/-^ F_2_ mice (Figure 6) was intriguing. However, while the number of F_2_ animals phenotyped here is sufficient to eliminate alternative receptor usage by IFNγ as the basis for IFNγR-independent Ab production, retroviral resistance is not a fully penetrant phenotype in either 129S1 or 129S7-*Ifngr*^*-/-*^ mice, and it appears that IFNγ-independent antiviral Ab production in 129S7 mice is a complex trait. Therefore, larger numbers of F_2_ mice would be required to draw any other conclusions regarding the genetic basis for these substrain differences.

While 129 substrains have the same genotypes at known MLV-resistance loci [(21, 23, 37, 38) and this report], our findings clearly indicate that there are unappreciated differences in their ability to mount protective antiviral immune responses. As such, it is unknown whether the 129P2/OlaHsd substrain utilized in many investigations on immune responses to FV infection would exhibit a similar phenotype to 129S1 mice when infected with RL-MLV, or whether they can produce IFNγ-independent antiviral immune responses. In fact, these substrains have been separated since the re-establishment of the 129 line in 1948 following the fire at Jackson Labs and have substantial genetic variation between them ((35, 36, 48) and Figure S1). The different phenotypes observed here between Steel substrains are not entirely surprising since the 129S1 lineage has been separated from the 129/SvEv lineage, from which the IFNγR^-/-^ 129S7 mice are derived, since 1969. Furthermore, additional crosses occurred in the 129/SvEv lineage prior to the introduction of the *Ifngr1* mutation in 129S7-derived AB1 ES cells [Figure S1 and (35, 48)], thereby increasing the genetic divergence of these substrains. Ideally, we would conduct experiments to confirm whether 129S7 and 129S1 mice inherit the same genetic mechanism controlling the production of anti-MLV Abs and differ only their ability to produce these responses in the absence of IFNγ. Unfortunately, such experiments cannot be conducted because wild-type 129S7 mice no longer exist. Collectively, our findings further emphasize the need for careful consideration of the genetic background of mutant and control mice and indicate that genetic differences between 129S substrains can have consequences well beyond affecting the efficiency of traditional gene targeting methods.

## Acknowledgements

We thank Dr. Tatyana Golovkina for support and consultation; S. Gingras and the transgenic and gene targeting (TGT) core for the generation of *Ifng*^*-/-*^ mice; Leigh Miller and Dr. John Alcorn for reagents and technical assistance for intracellular IFNγ staining; and members of the Kane, Golovkina, and Dr. Alexander Chervonsky laboratories for helpful discussion. This work was supported by a Pilot Award from the RK Mellon Institute for Pediatric Research (to M.K.). R.Z. is supported by T32 AI049820. The content is solely the responsibility of the authors.

## Author contributions

Conception and design: M.K.; Acquisition of data: R.Z.Z., V.M., L.R., and M.K.; Analysis and interpretation of data: R.Z.Z. and M.K.; Drafting the article: M.K.; Revising the article: M.K., R.Z.Z., and V.M..

## Materials and Methods

### Mice

129S1/SvImJ (129S1 stock #002448) and BALB/cByJ (stock #001026) were purchased from the Jackson Laboratory. 129-*Ifngr1*^*tm1Agt*^/J [(48) also known as G129] mice were purchased from the Jackson Laboratory (via cryo recovery, stock #002702) and bred and maintained at the University of Pittsburgh. IFNγR^-/-^ 129 mice were generated using AB1 ES cells derived from 129S7/SvEvBrd-*Hprt*^*b-m2*^ (129S7) mice, and chimeric founder males were crossed to an unknown 129/SvEv substrain (although 129S8 mice are commonly referred to as 129/SvEv and may have been used for these crosses) (48) for simplicity, we refer to these as 129S7-*Ifngr*^*-/-*^ mice (Figure S1). C.129S7(B6)-*Ifngr1*^*tm1Agt*^/J (IFNγR^-/-^ ByJ) were purchased from the Jackson Laboratory (via cryo recovery, stock #002286) and bred and maintained at the University of Pittsburgh. Mice of both sexes were used in equal ratios for all experiments.

*Ifng*-deficient 129S1 mice were generated using CRISPR/Cas9 technology. The target sequences (5′ guide sequence: 5′-GCTGTTTCCCTGCGTAGTTT-3′; 3′ guide sequence: 5′-TAGAGGCTAACCAGAGCCGA-3′) were previously validated for the generation of conditional knockout strains on the C57BL/6 background (43). Two male founders with complete deletions of exons 2 and 3 (removing sequences coding for amino acids 38-120) were selected (Figure S3A) and crossed to 129S1 females. Potential off-target sites with fewer than three mismatches identified by the Cas-OFFinder algorithm (49) were sequenced in N_1_ and N_2_ mice, and mice with no mutations at these sites were intercrossed to produce homozygous *Ifng*-deficient mice. Both founder lines express a truncated IFNγ protein [lacking four of six alpha helices (50)] that is not biologically active (Figure S3B-C).

Mice were genotyped from tail biopsies using real time PCR with specific probes designed for each gene (Transnetyx, Cordova, TN) or by flow cytometry (see below). Mice of both sexes were used in equal numbers for each experiment. All animal experiments were performed in the American Association for the Accreditation of Laboratory Animal Care-accredited, specific-pathogen-free facility Division of Laboratory Animal Resources, University of Pittsburgh School of Medicine. Animal protocols were reviewed and approved by the Institutional Animal Care and Use Committee at The University of Pittsburgh.

### Flow cytometry

For MHC genotyping of [(129S1xByJ)xByJ]N_2_ mice, blood was collected in sodium heparin collection vials (SAI Infusion Technologies), and peripheral blood lymphocytes were isolated by overlaying with Ficoll-Paque (Cytivia) and spinning at 11,000xg for 4min at 25^°^C. Lymphocytes were stained with antibodies against Class I [brilliant blue 700 (BB700)-conjugated anti-H-2Db (BD Biosciences) and phycoerythrin (PE)-conjugated anti-H-2Dd (BD Biosciences)] or Class II [peridinin chlorophyll (PerCP)-eFluor 710-conjugated anti-H2-Ab1 (I-Ab, Invitrogen) and PE-conjugated anti-H2-Ad1 (I-Ad, BD Biosciences)] MHC and analyzed on an Attune NxT cytometer (Life Technologies).

### MLV infection

Rauscher-like MuLV (RL-MuLV), a mixture consisting of NB-tropic ecotropic and mink lung cell focus-forming viruses, was described previously (69) and was provided by T. Golovkina, University of Chicago. The virus was propagated in SC-1 embryonic mouse fibroblasts (ATCC). Ecotropic (Eco) viral titers were determined by an infectious center assay (70). Experimental mice were injected i.p. with 2 ×10^4^ Eco PFUs at 5-8 weeks of age and screened for anti-virus antibodies and plaque forming units 8-10 weeks later.

### ELISA

To detect MuLV Abs in mouse sera, an enzyme-linked immunosorbent assay (ELISA) was performed as previously described (6, 15). Virions isolated from RL-MLV infected SC-1 cells were treated with 0.1% Triton X-100 and bound to plastic in borate-buffered saline overnight, followed by incubation with mouse serum samples at 4^°^C for 90 minutes. All sera were used at 2 × 10^−2^ dilutions. Mouse IgG2a-specific, as well as total IgG-specific secondary antibodies coupled to horseradish peroxidase (HRP) (Jackson ImmunoResearch) were used to detect anti-virus antibodies. Ovalbumin (2%) was used as a blocking reagent. Backgrounds obtained from incubation with secondary antibodies alone were subtracted from the values obtained from sera of infected mice.

### Statistical analyses

Statistical significance was determined using GraphPad software (one way ANOVA or unpaired t test).

## Supplemental Materials and Methods

### Splenocyte isolation for stimulation

The spleens of mice were aseptically isolated, trimmed of all excess tissue and placed in sterile phosphate-buffered saline (PBS, Corning). Cell suspensions were erythrocyte-depleted by incubation in 450µl sterile distilled water (Gibco) for 10 seconds before the addition of 50µl of 10X PBS (Gibco) and 4mL of 1X PBS. Cell suspensions were pelleted at 300x*g* for 5 min at 25°C. Splenocytes were resuspended in RPMI 1640 Medium (Gibco), supplemented with 10% heat-inactivated fetal calf serum (FCS, Gibco) and 50µg/ml gentamicin (Gibco).

### Flow cytometry

Splenocytes (2 × 10^6^) were stimulated with 50ng/ml phorbol 12-myristate 13-acetate (PMA, Sigma-Aldrich) and 750ng/ml ionomycin (Sigma-Aldrich) at 37°C with 5% CO_2_ for six hours. For the last five hours, 1µg/ml GolgiPlug™ Protein Transport Inhibitor (BD Biosciences) was added to block cytokine secretion. Cells were surface stained with antibodies against AlexaFlour 532-conjugated CD45 (clone 30-F11, Invitrogen) in PBS supplemented with 0.5% BSA, 0.1% sodium azide, 3mM egtazic acid (EGTA), and 20ug/mL DNAse1. Cells were then fixed and permeabilized with eBioscience™ Foxp3/Transcription Factor Staining Buffer Set (Invitrogen) and stained intracellularly with phycoerythrin (PE)-conjugated anti-IFNγ (clone XMG1.2 RUO, BD Bioscience) in permeabilization buffer. Cells were stained for viability with Zombie NIR™ Fixable Viability Kit (BioLegend) in PBS and Mouse Fc Block™ (BD Biosciences). Samples were analyzed on a Cytek® Aurora (Cytek Biosciences).

### IFNγ detection assay

IFNγ detection assay for biologically active mouse IFNγ was performed using B16-Blue™ IFNγ reporter cells (InvivoGen) engineered to produce secreted alkaline phosphatase (SEAP) in response to IFNγ stimulation. Cells were cultured following the manufacturer’s protocol in Dulbecco’s Modified Eagles Medium (DMEM, Gibco) supplemented with 10% FCS, 50µg/ml gentamicin, 100 µg/ml Normocin™ (InvivoGen), and 100 µg/ml Zeocin® (InvivoGen).

Splenocytes (2 × 10^6^) were stimulated with 50ng/ml PMA and 750ng/ml ionomycin in 200µl of cell culture medium at 37°C/5% CO_2_ for 16 hours. After stimulation, 20µl of supernatant were added per well in a 96-well flat bottom plate. To generate a standard curve, 20µl recombinant murine IFNγ (MilliporeSigma™ 40732020UG) was added at a starting concentration of 100ng/ml and serially diluted (five-fold dilutions). After addition of samples, 7.2 × 10^4^ B16-Blue™ IFNγ cells were added in 180µl of cell culture medium for a final working volume of 200µl. Samples were incubated at 37°C/5% CO_2_ for 24 hours.

For QUANTI-Blue detection of SEAP, 20µl of supernatant was added to a 96-well flat bottom plate followed by addition of 180μl of QUANTI-Blue Solution™ (InvivoGen) and incubated at 37°C for 24 hours. SEAP levels were determined using a Synergy H1 microplate reader (BioTek) by reading the optical density (OD) at 650nm. Background was subtracted with a negative control of QUANTI-Blue Solution™ incubated with 20µl of complete cell culture media. Levels of IFNγ in splenocyte supernatants were determined based on a standard curve for recombinant murine IFNγ.

### Immunoblot Analysis

RL-MLV virions were isolated from supernatants of infected SC-1 cells via centrifugation at 95,000x*g*. MLV virions were lysed in NuPage LDS sample buffer (Novex), separated by electrophoresis on NuPage 4–12% Bis-Tris gels (Invitrogen) and blotted onto polyvinylidene fluoride (PDVF, BioRad Laboratories). Membranes were incubated with sera at 5 × 10^−3^ dilution. For the second step, membranes were incubated with goat anti-mouse-IgG2a-specific antibodies coupled to HRP (Jackson ImmunoResearch). Blots were developed with SuperSignal™ West Pico PLUS Chemiluminescent Substrate (Thermo Scientific) and imaged on the ChemiDoc Imaging System (Bio-Rad).

**Table S1.**
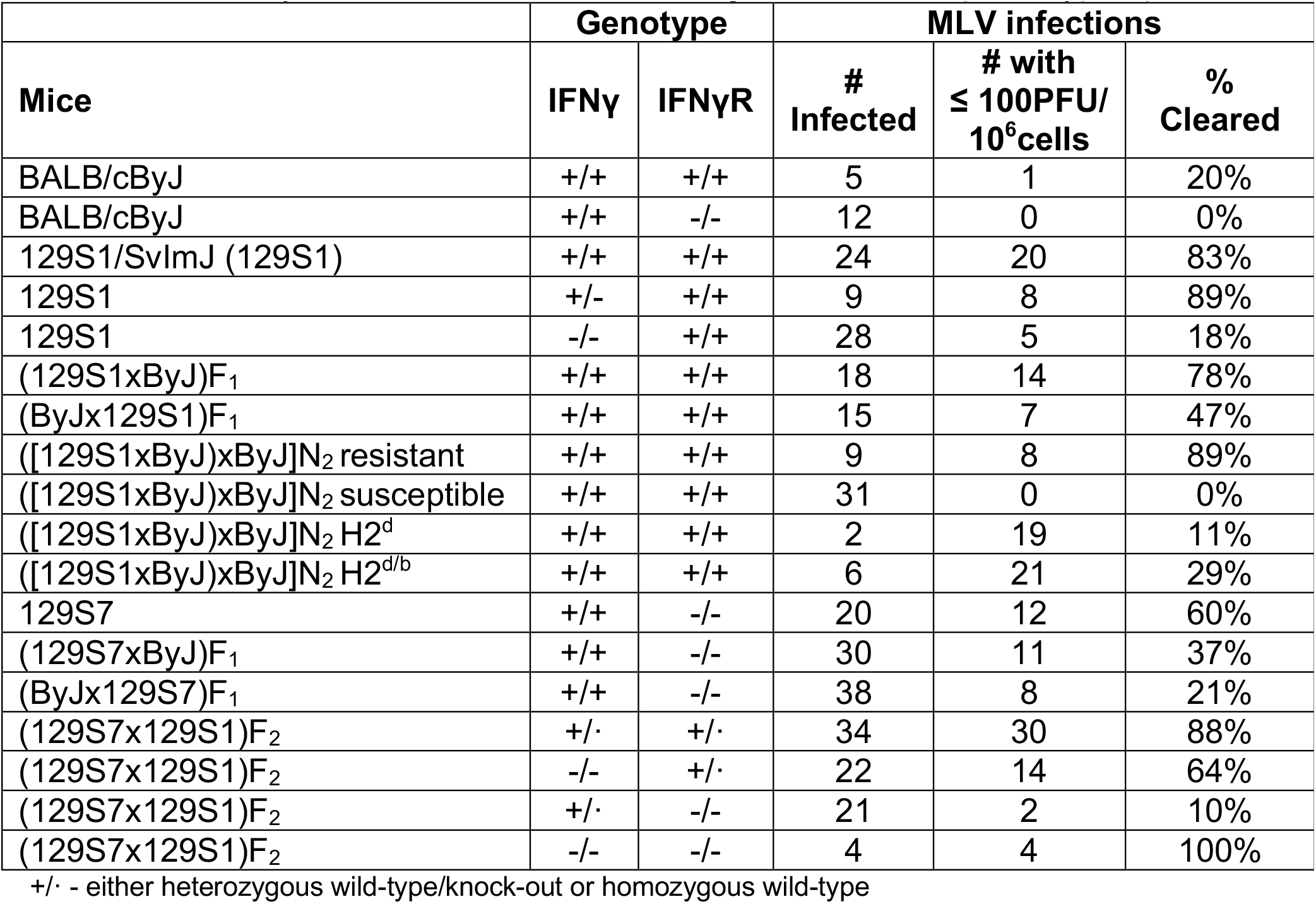
Summary of mice utilized in this investigation and their phenotype upon MLV infection

**Figure S1.**
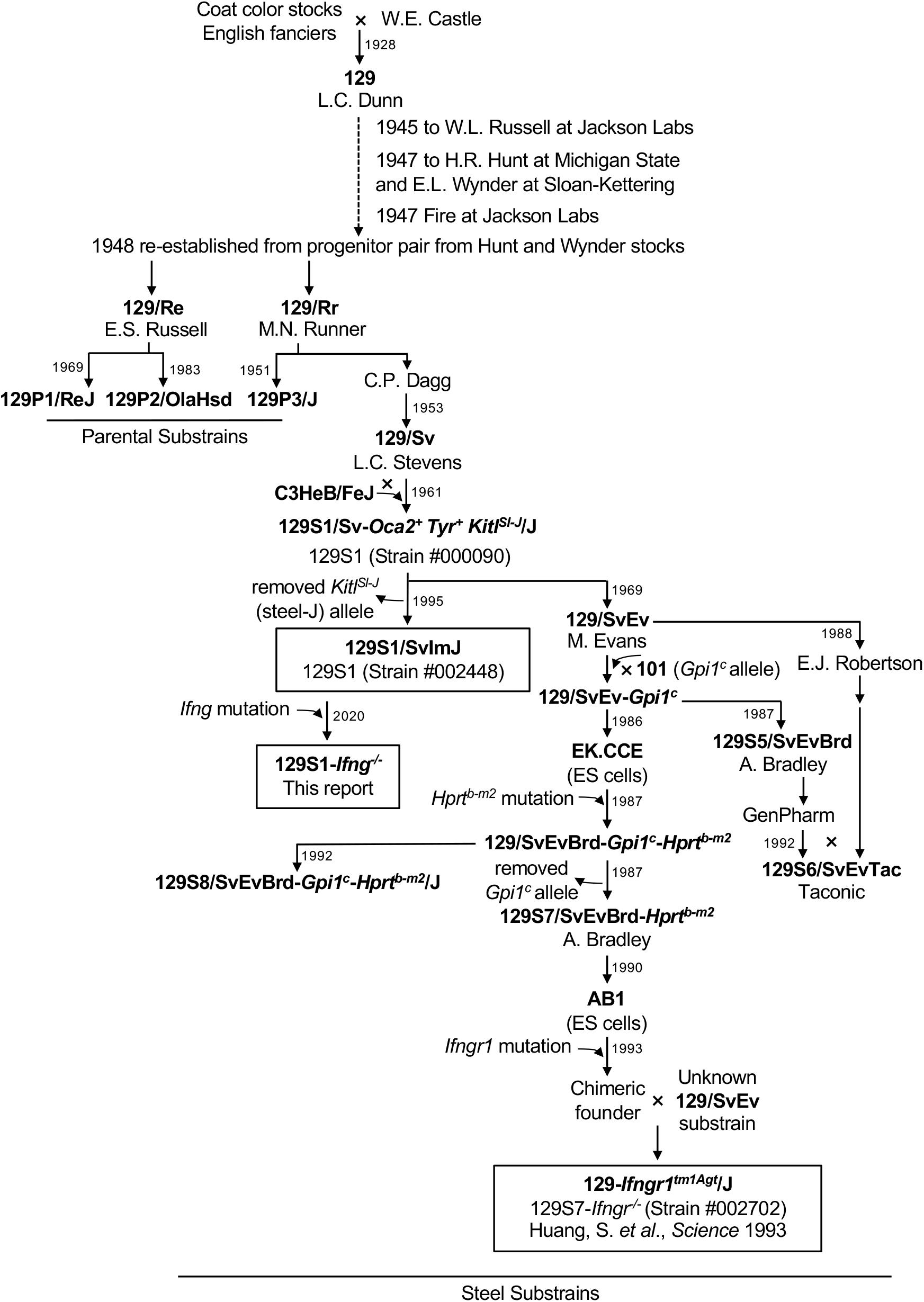
History of selected 129 sublines. Schematic of the genealogy of 129S1 and 129S7-*Ifngr*^-/-^ mice. Adapted from (35).

**Figure S2.**
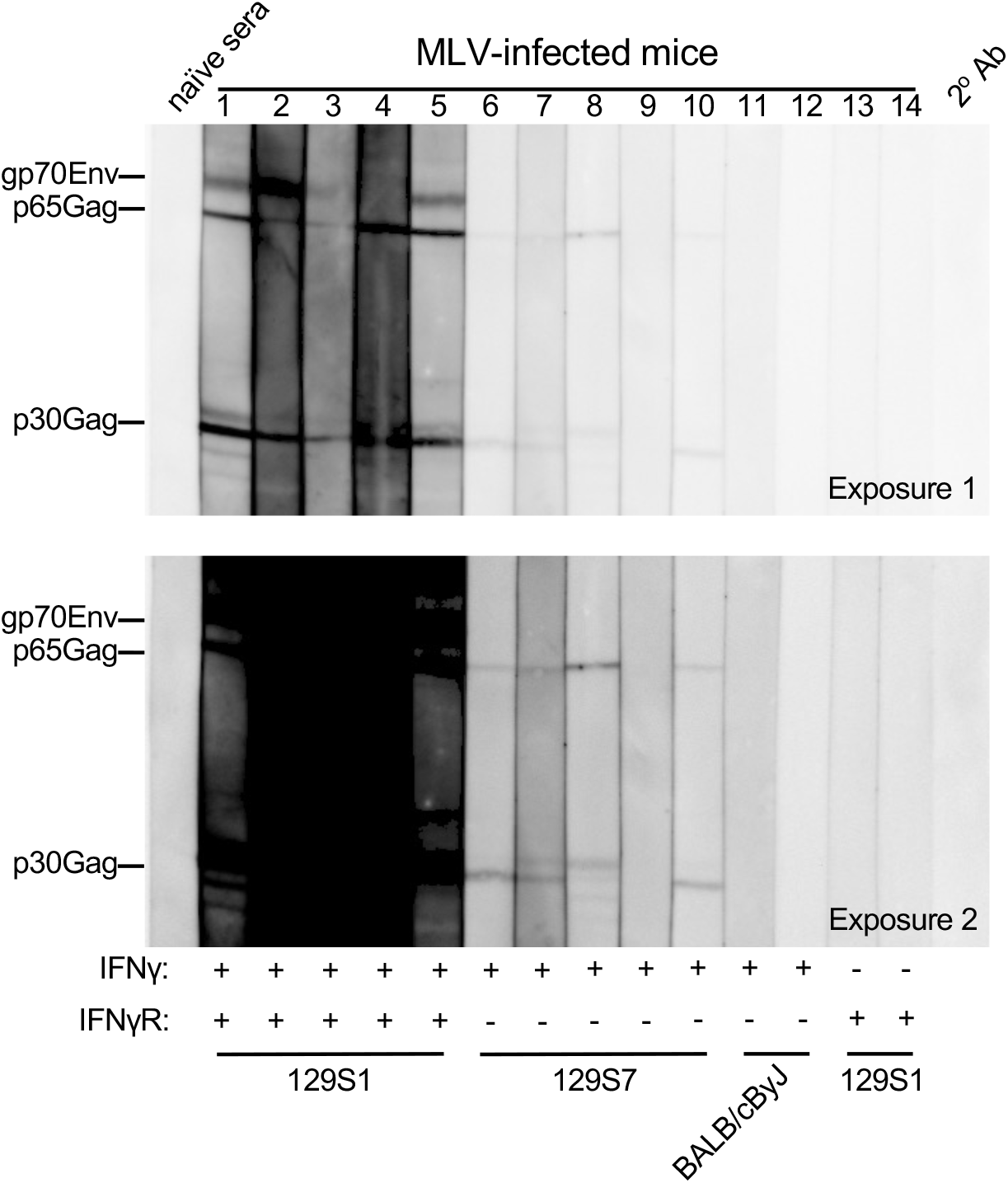
Specificity of antiviral Abs. Sera from 129S and BALB/cByJ mice of the indicated genotypes were tested for reactivity against MLV virion proteins by immunoblot 10 weeks post infection. Goat anti-mouse IgG2a-specific Abs coupled to HRP were used at the second step. Numbers correspond to individual mice. Two different exposures are shown.

**Figure S3.**
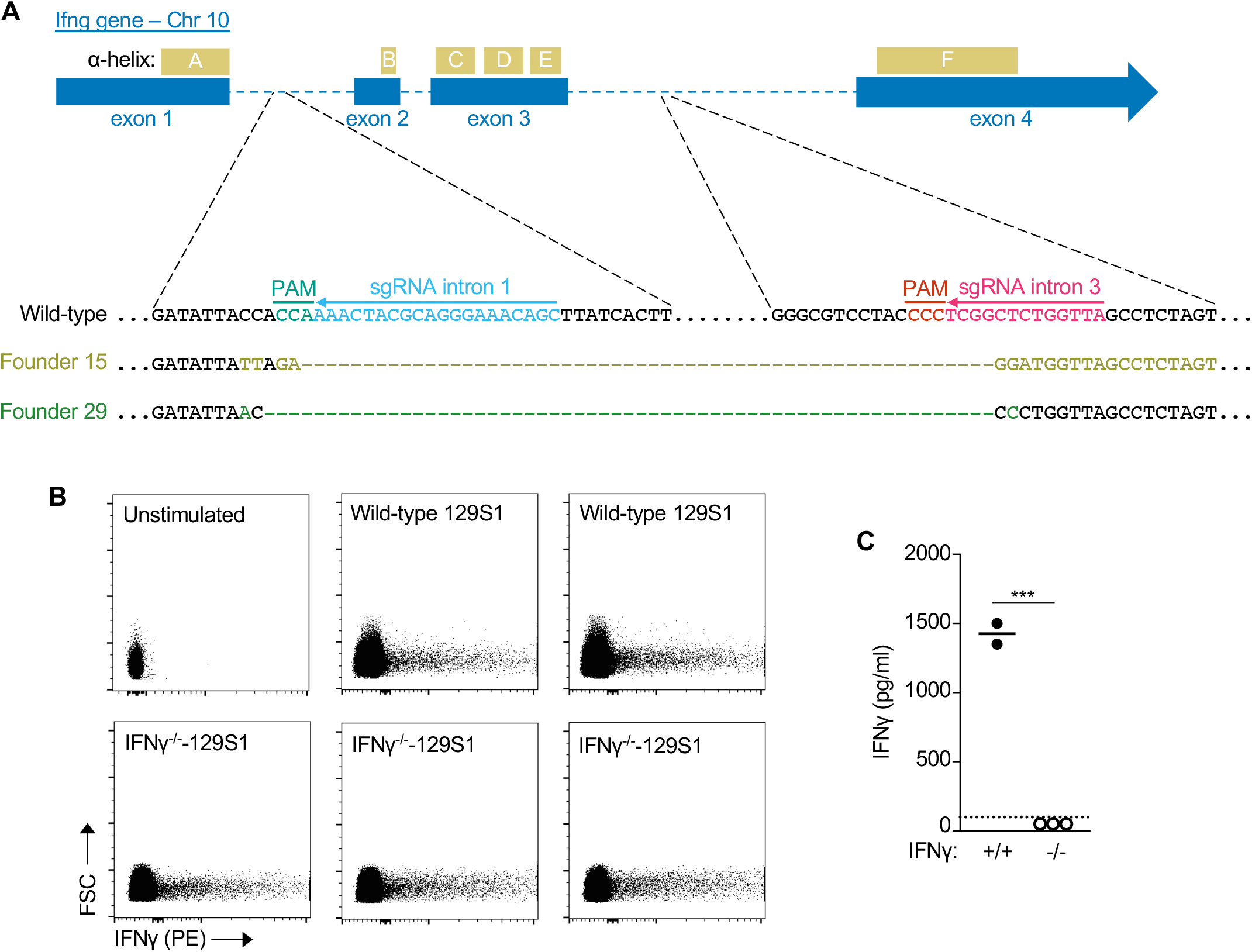
Generation of IFNγ-deficient 129S1 mice. **A)** Diagram of the *Ifng* locus, with sgRNA targeting sited in introns 1 and 3 indicated. The six *α*-helices based on the human IFNγ structure (REF) are indicated above the exons. Sequence of wild-type mice at targeted sites, with sgRNA and PAM sites indicated. Dots indicate sequences flanking those shown in detail. Sequences of the locus in selected founder mice with mismatches to wild-type sequence highlighted and deleted sequence indicated by dashes. Both founder lines lack exons 2 and 3, which encode for amino acids 38-120 (helices B through E). **B)** Truncated IFNγ is expressed in IFNγ^-/-^-129S1 mice. Flow-cytometric plots showing intracellular IFNγ in unstimulated and PMA/ionomycin-stimulated CD45^+^ splenocytes from wild-type 129S1 and IFNγ^-/-^-129S1 (Founder 29) mice. **C)** The truncated IFNγ expressed in 129S1-*Ifng*^-/-^ mice is not biologically active. Detection of IFNγ in supernatants from PMA/ionomycin-stimulated splenocytes from wild-type 129S1 and IFNγ^-/-^-129S1 (Founder 29) mice by B16-Blue™ IFNγ cells. Dashed line indicates the limit of detection (100pg/ml).

